## Supplementary Figures, Tables & Pipelines for "Both Genome Instability and Replicative Senescence Stem from the Shortest Telomere in Telomerase-Negative Cells"

Contents:

Supplementary Fig. 1-7
Supplementary Tables 1-13
Supplementary Pipeline 1
Supplementary Pipeline 2
References

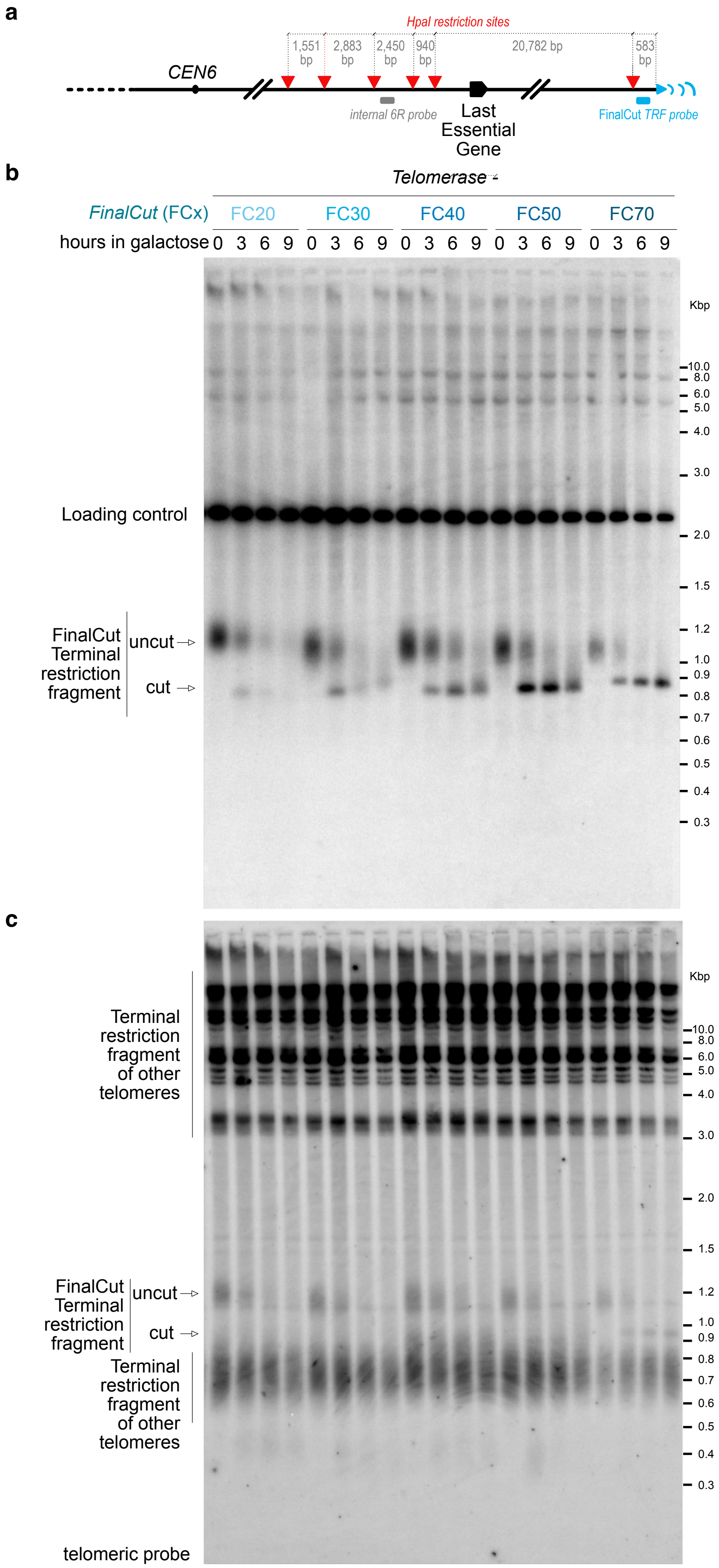
a
HpaI restriction sites
1,551 bp
2,883 bp
2,450 bp
940 bp
20,782 bp
583 bp
CEN6
internal 6R probe
Last Essential Gene
FinalCut TRF probe
b
Telomerase -
FinalCut (FCx)
FC20
FC30
FC40
FC50
FC70
hours in galactose 0 3 6 9 0 3 6 9 0 3 6 9 0 3 6 9 0 3 6 9
Kbp
10.0
8.0
6.0
5.0
4.0
3.0
2.0
1.5
1.2
1.0
0.9
0.8
0.7
0.6
0.5
0.4
0.3
Loading control
FinalCut Terminal restriction fragment
uncut
cut
c
Kbp
Terminal restriction fragment of other telomeres
FinalCut Terminal restriction fragment
uncut
cut
Terminal restriction fragment of other telomeres
telomeric probe

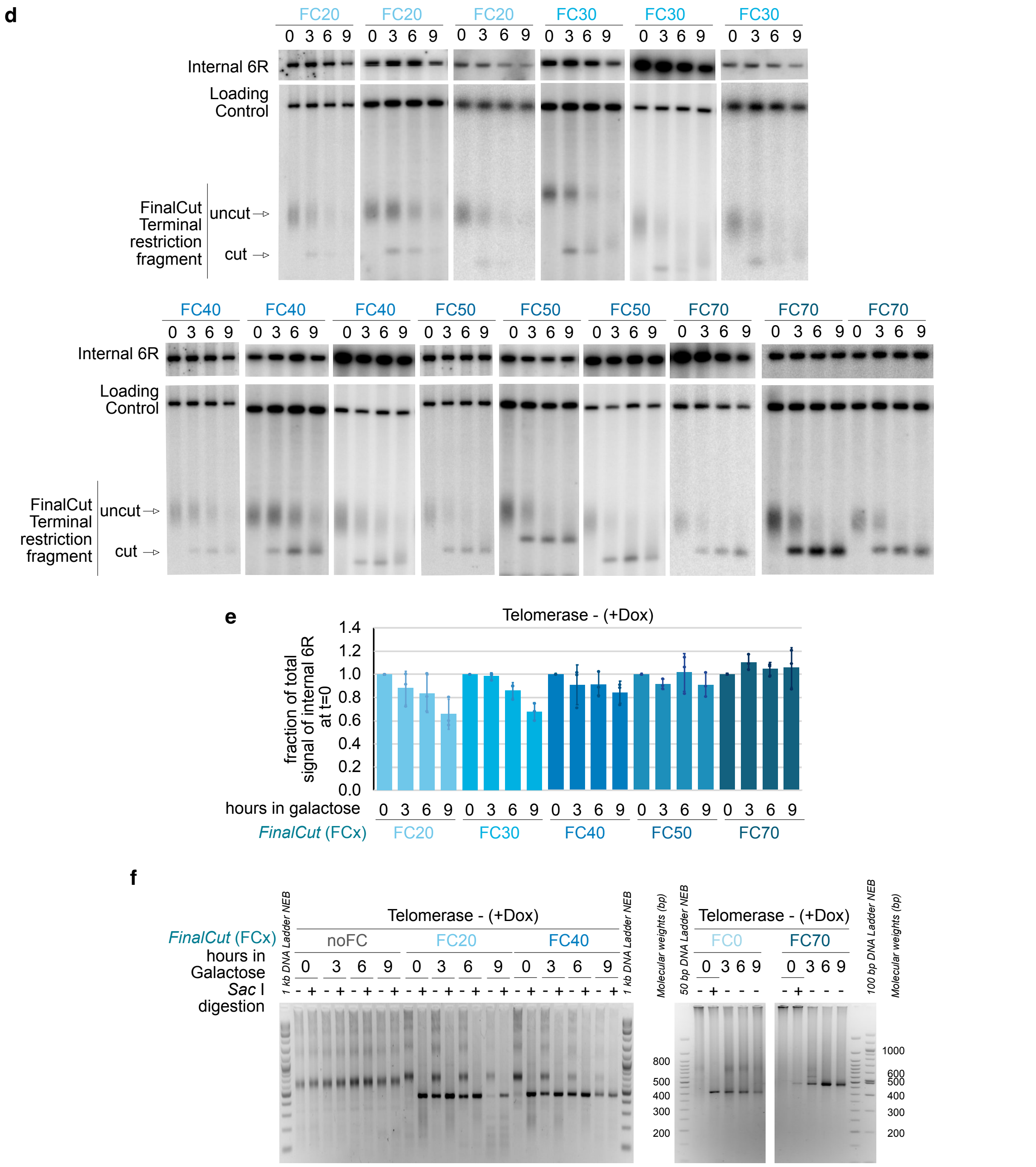


***Supplementary Fig. 1: Related to Fig.1.***

***a,*** *Restriction map of the end of chromosome 6R arm (not to scale).*

***b-c,*** *The Southern blot shown in Figure 1d shown in full (b) was rehybridized with an oligo probe targeting telomeric repeats (c).*

***d,*** *Triplicate individual Southern blots of TRF of* Hpa *I digested genomic DNA, prepared at indicated time points from strains with indicated* FinalCut *telomere lengths in the absence of telomerase, probed with a subtelomeric 6R to detect FinalCut, an internal locus in chromosome 11 (Loading control), and an internal region of chromosome 6 R arm 23 kb away from FinalCut, used for quantifications shown in Supplementary Fig. 1e.*

***e,*** *Stability of the right arm of chromosome 6 at 23 kb from the FC telomere as quantified from Southern Blots shown in d. N=3 independent experiments. Error bars correspond to SD.*

***f,*** *Efficiency of Cas9-dependent* FinalCut *cleavage was also qualitatively verified by TeloPCR of* TEL6R*. Here, representative result of TEL6R-specific telo-PCR with genomic DNA of indicated strains incubated in doxycycline and galactose for indicated hours. If indicated, PCR product was digested with* Sac *I restriction enzyme at 37°C for 1 hour.* Sac *I cleaves intact ttDNA (see Fig. 1a). TeloPCR products were migrated in 2.5% agarose gels.*
18 *TEL6R-specific primer oT883 binds 6R subtelomere at 370 bp from the sub-telomere-telomere junction.*

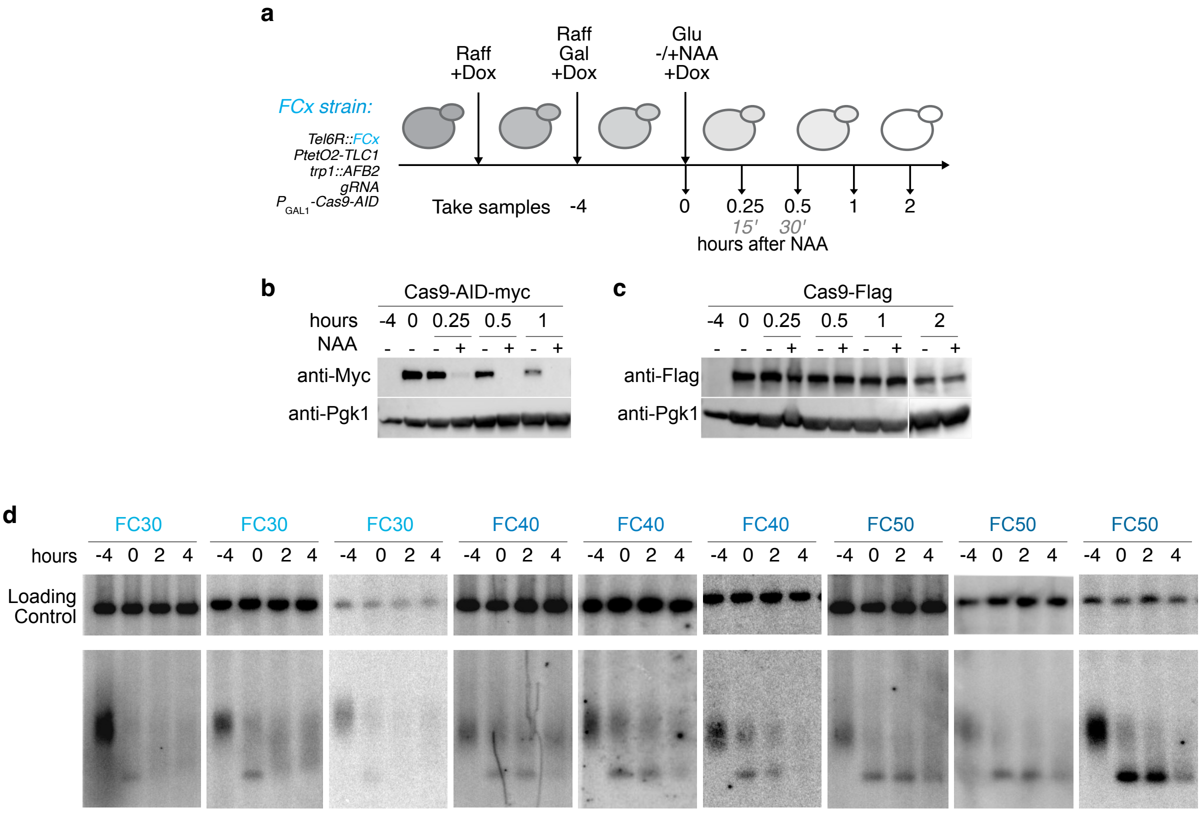


**Supplementary Fig. 2: Related to Fig. 2.**

***a,*** *Experimental design for testing Cas9-AID degradation.*

***b-c,*** *Representative western blot of Cas9-AID fused to a Myc-tag (b) or a Cas9 without AID fused to a Flag-tag (c) in a FC70 strain. An anti-Pgk1 serves as loading control.*

***d,*** *Individual Southern blots of TRF of* Hpa *I digested genomic DNA, prepared at indicated time points from strains with indicated* FinalCut *telomere lengths in the absence of telomerase, probed with the subtelomere 6R and an internal locus in chromosome 11 (Loading control), used for quantifications shown in Fig. 2c.*

19

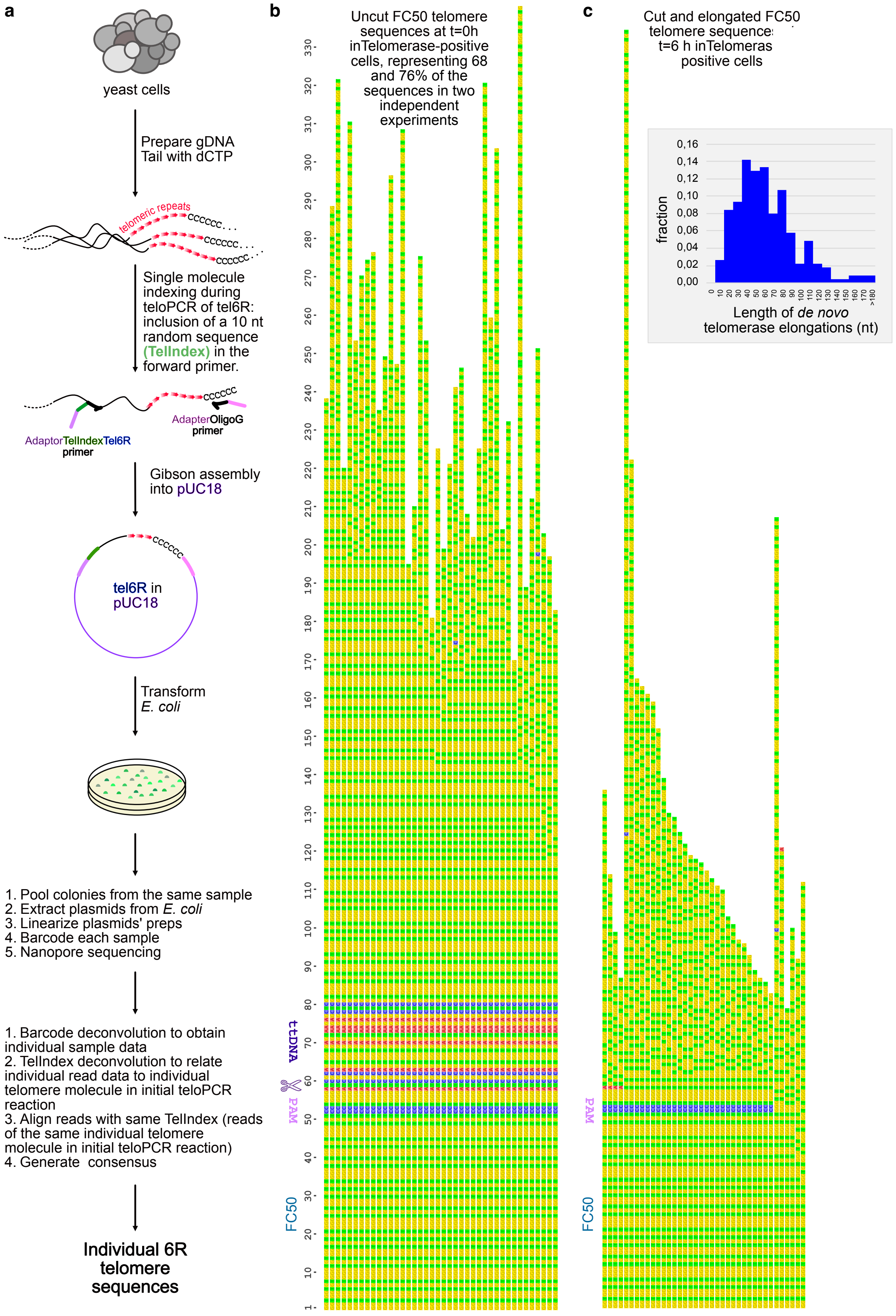
a
yeast cells
Prepare gDNA
Tail with dCTP
telomeric repeats
CCCCCC...
Single molecule indexing during teloPCR of tel6R: inclusion of a 10 nt random sequence (TelIndex) in the forward primer.
AdaptorTelIndexTel6R primer
AdapterOligoG primer
Gibson assembly into pUC18
tel6R in pUC18
Transform E. coli
1. Pool colonies from the same sample
2. Extract plasmids from E. coli
3. Linearize plasmids' preps
4. Barcode each sample
5. Nanopore sequencing
1. Barcode deconvolution to obtain individual sample data
2. TelIndex deconvolution to relate individual read data to individual telomere molecule in initial teloPCR reaction
3. Align reads with same TelIndex (reads of the same individual telomere molecule in initial teloPCR reaction)
4. Generate consensus
Individual 6R telomere sequences
b
Uncut FC50 telomere sequences at t=0h inTelomerase-positive cells, representing 68 and 76% of the sequences in two independent experiments
t:tDNA
PAM
FC50
c
Cut and elongated FC50 telomere sequence t=6 h inTelomeras positive cells
fraction
Length of de novo telomerase elongations (nt)
PAM
FC50

**d**

FC50 telomere sequences at t=4 h inTelomerase-negative cells

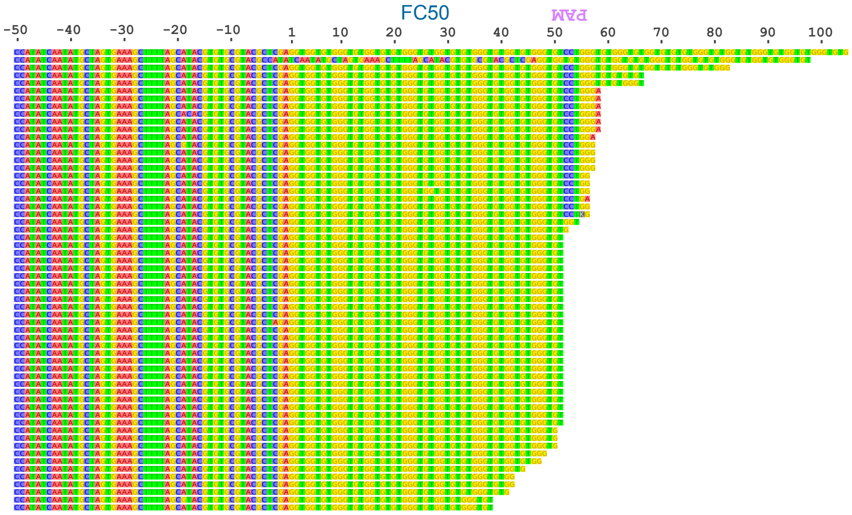


**e**

FC50 telomere sequences at t=2 h inTelomerase-negative cells

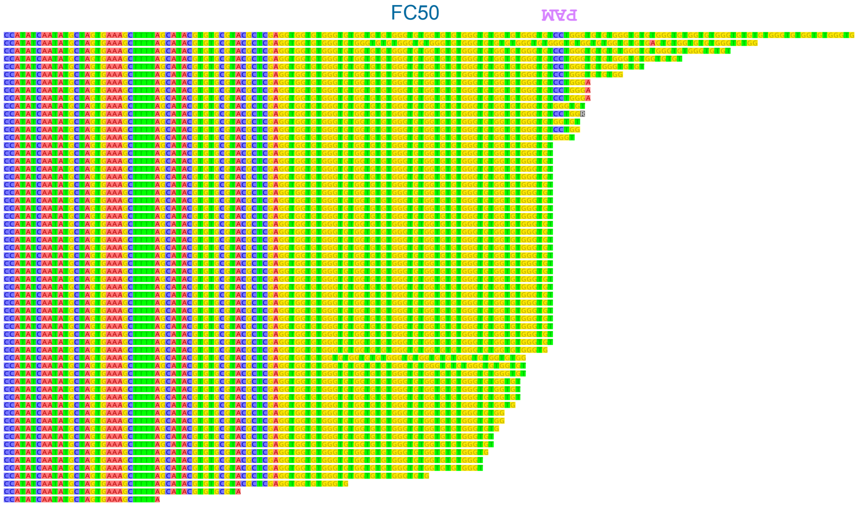


**f**

FC50 telomere sequences at t=4 h inTelomerase-negative cells

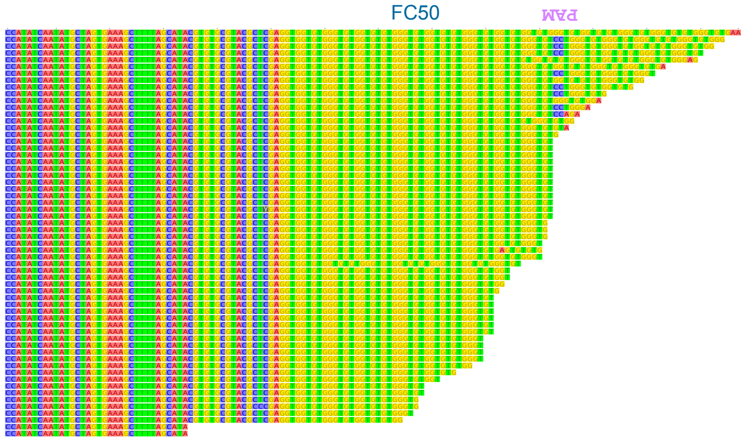

**g**

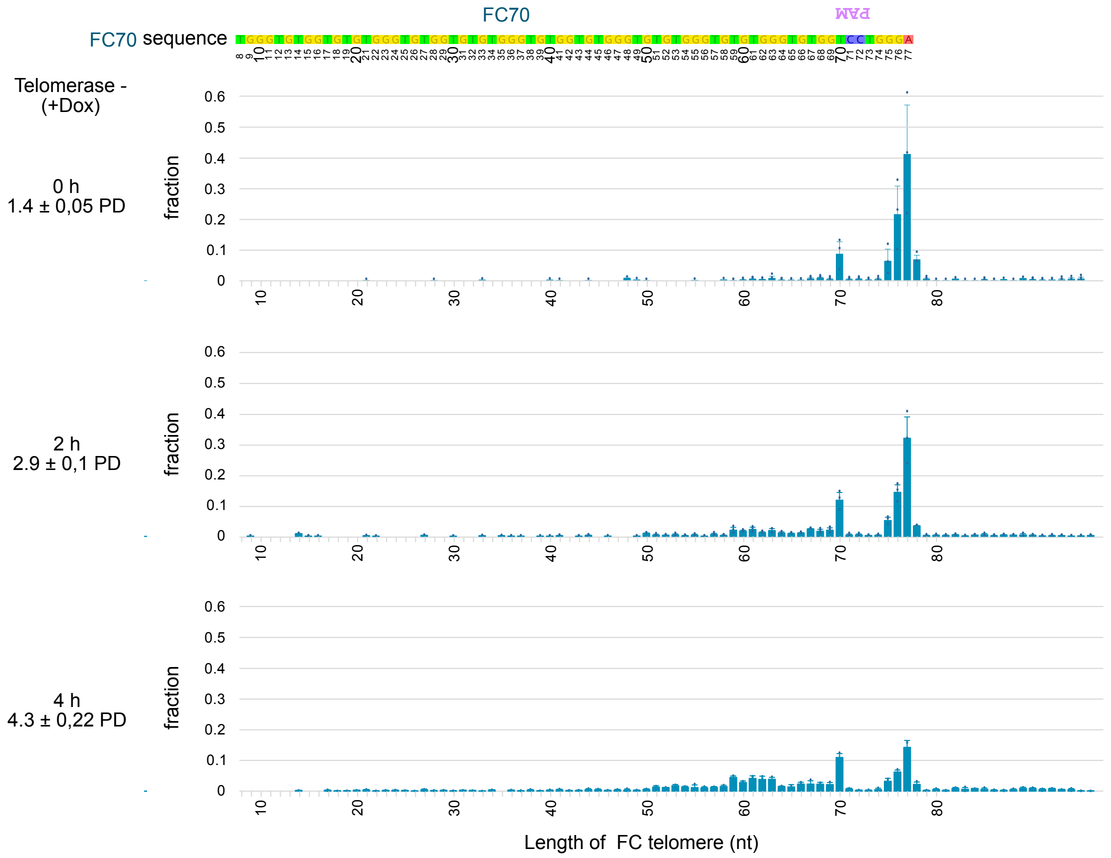


***Supplementary Fig. 3: Related to Fig. 3.***

***a,*** *Protocol of sample preparation to sequence of hundreds of individual telomeres with high accuracy.*

***b-f,*** *Examples of sequences of individual FC50 telomeres before (b) or after indicated times in Galactose induction of Cas9 during 4 h and subsequent shift into glucose and NAA in the absence (c) or presence of doxycycline (d-e). 20 consensus sequences of 2 (b), 19 of 2 (c) or 20 of 3 (d-e) independent experiments are shown for each time point. Sequences were edited for single mismatches sequencing errors. Inset in (c) corresponds to the distribution of telomerase-dependent re-elongations evaluated from sequence divergence.*

***g,*** *Distribution of individual telomere lengths with time after Cas9 cut as in Fig.3f using FC70 strain in the absence of telomerase (+Dox) for indicated range of telomere lengths/Magnified view of Fig.3e. Dots represent the average of each of 3 independent experiment in which N=1119, 1109, 764 for t=0 h; N=819, 831, 1416 for t=2 h; N=1102, 1018, 745 for t=4 h individual telomere sequences, as indicated in the legend of Fig. 3e. Error bars correspond to SD from the average of 3 independent experiments.*

22

23

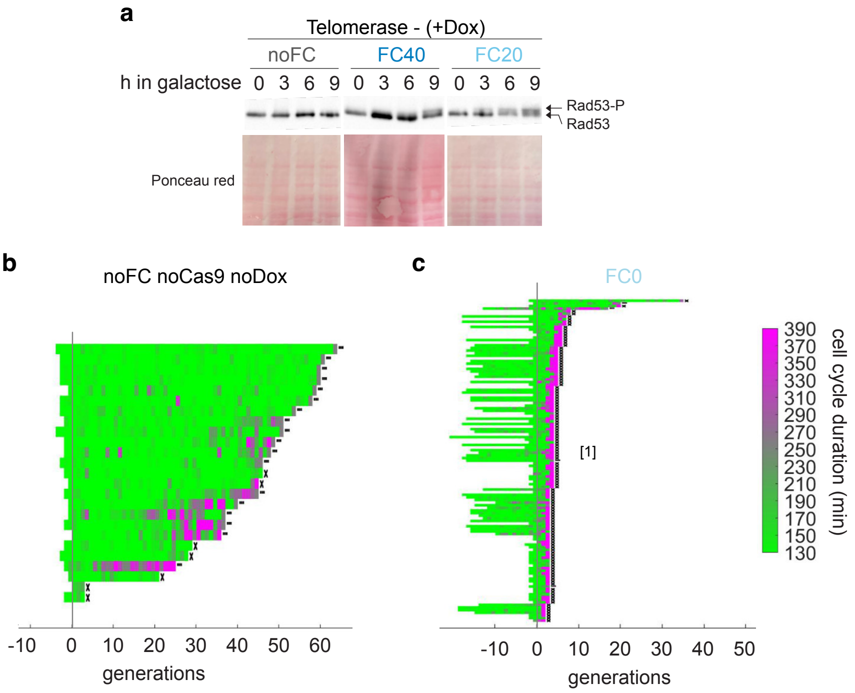


***Supplementary Fig. 4: Related to Fig. 4.***

***a,*** *Indicated FinalCut strains were cultured in media containing doxycyclin and carbon source was changed to galactose for indicated time. Proteins were extracted and Rad53 phosphorylation was monitored by Western blot using an anti-Rad53 antibody. Ponceau red staining is shown as loading control.*

***b-c,*** *Microfluidics positive and negative controls. Display of consecutive cell cycle durations of lineages of indicated strains. Experimental scheme and legends as in Fig. 4c. For b, recording was stopped at 5 days post injection into the microfluidics circuit (note many lineages haven't died at the end of the experiment (labelled with ellipsis (...)).*

24

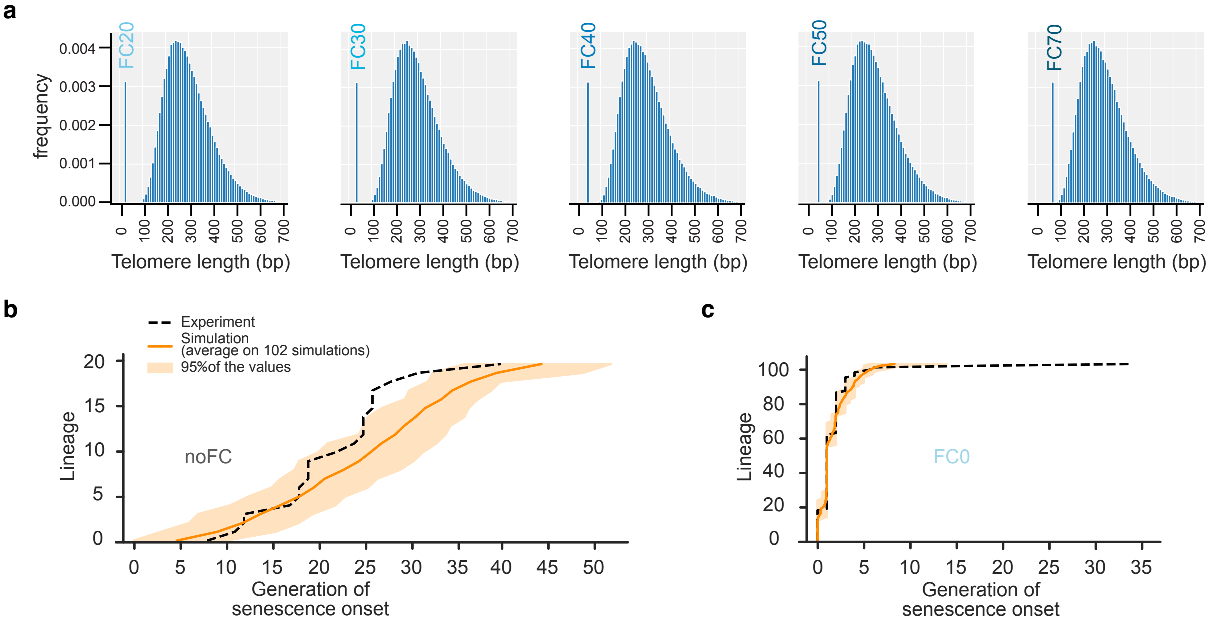


***Supplementary Fig. 5: Related to Fig. 4.***

***a,*** *Telomere length distribution in cells that experienced a cut across 50 000 simulated lineages, with the mathematical model set for indicated* FinalCut *conditions. Values above 700 bp were truncated for clarity.*

***b-c,*** *Lifespan of individual cell lineages entering senescence upon telomerase inactivation, as assessed by microfluidics in Supplementary Fig. 4b-c, and 102 simulations of the same number of individual cell lineages with the mathematical model set for indicated* FinalCut *conditions, ordered by lifespan.*

25

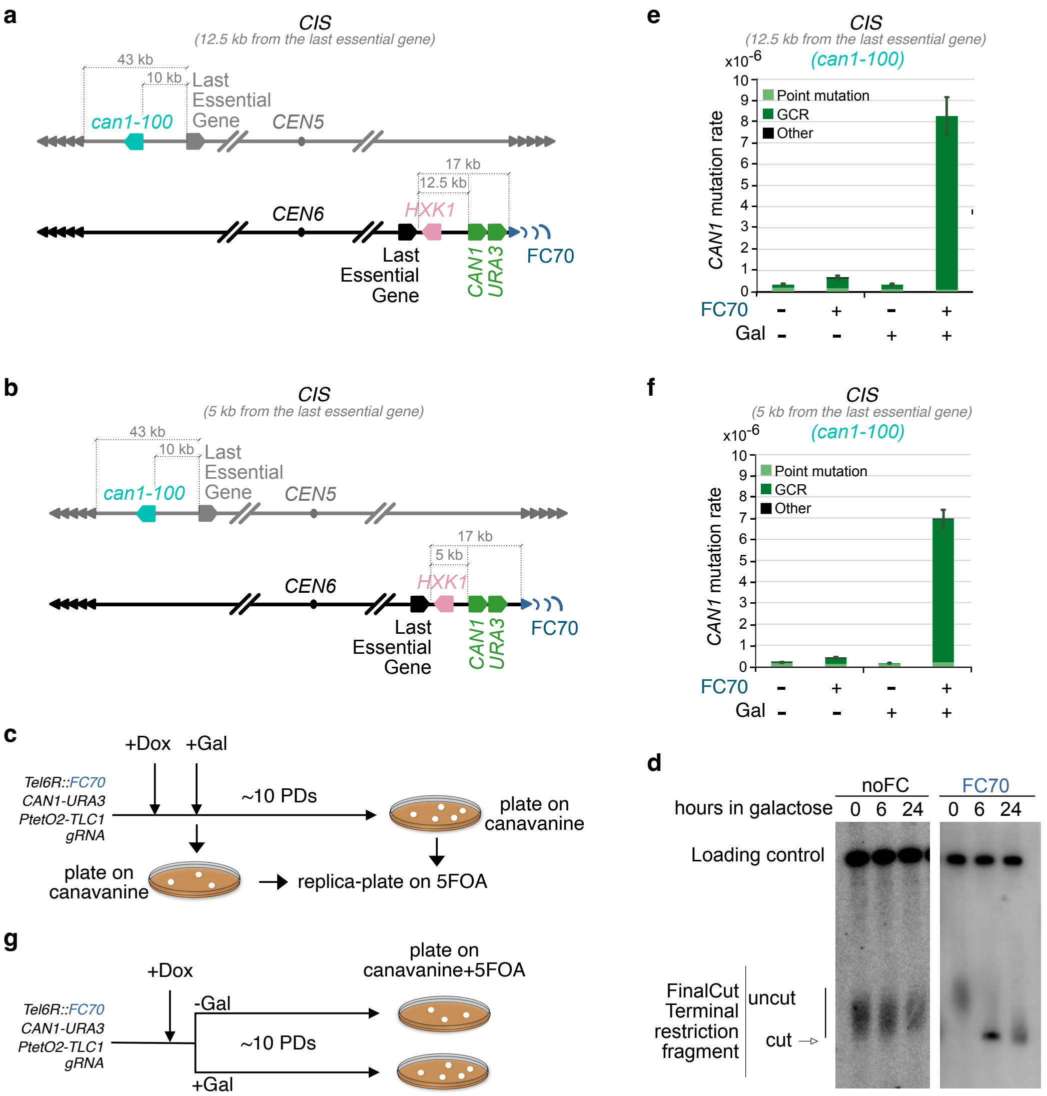


**Supplementary Fig. 6: Related to Fig. 6.**

***a-b,*** *Relevant genomic features of strains used to quantify genomic instability at 12.5 kb (a: yT1696) or 5 kb (b: yT1698) from the last essential gene and in cis of* FC70 *. Legends as in Fig. 6a-b.*

***c,*** *Experimental protocol for the fluctuation assay in cells harboring a* FC70 *telomere to determine CAN1 mutation rate. Multiple cultures are grown in parallel in which telomerase is inactivated by doxycycline addition, followed by carbon source switch to galactose to induce* FC70*. Then, a portion of each culture is plated at t=0 h and t=24 h after Cas9 induction on canavanine-containing glucose media lacking doxycycline. This allows for Cas9 shut off, telomerase re-expression to stabilize the genome and selection of mutants at CAN1 locus.* $Can^R$ *colonies number distribution is used to estimate mutation rate according to fluctuation assay mathematical models.* $Can^R$ *cells are then replica-plated on plates containing 5-FOA to score the fraction of GCRs or point mutations.*

***d,*** *Southern blot of strains carrying indicated constructs at TEL6R incubated for indicated hours in galactose with probe targeting the* FinalCut *TRF and loading control. Strains used: yT576 and yT1612.*

***e-f,*** *Mutation rate of* CAN1 *gene, located either 12.5 kb (d) or 5 kb (e) from the last essential gene and in* cis *of* noFC *or* FC70 *in* can1-100 *cells, calculated from cells plated at 0 h and 24 h after Cas9 induction (see Supplementary Table ). Data are represented as the mutation rate ±95% confidence interval. Mutation type distribution of* $Can^R$ *colonies obtained at 0 h and 24 h after Cas9 induction is indicated. Others correspond to a very small fraction of colonies [*$Can^R$ *5-FOA*$^R$ *ura*$^-$*]. See Table S9 for statistical details. Strain used in d: yT1696 (*FC70*) and yT1710 (*noFC*). Strain used in e: yT1698 (*FC70*) and yT1712 (*noFC*).*

***g,*** *Experimental protocol for the fluctuation assay to assess* CAN1-URA3 *mutation rate (GCRs). Multiple cultures are grown in parallel in which telomerase is inactivated by doxycycline addition, followed by carbon source switch to galactose to induce* FC70 *or not. Uppon ~10 PDs, the cultures are plated on canavanine and 5-FOA-containing glucose media lacking doxycycline. This allows for Cas9 shut off, telomerase re-expression to stabilize the genome and selection of mutants at both* CAN1 *and* URA3 *locus.* $Can^R$ *5FOA*$^R$ *ura3*$^-$ *colonies number distribution is used to estimate GCR rate according to fluctuation assay mathematical models.*

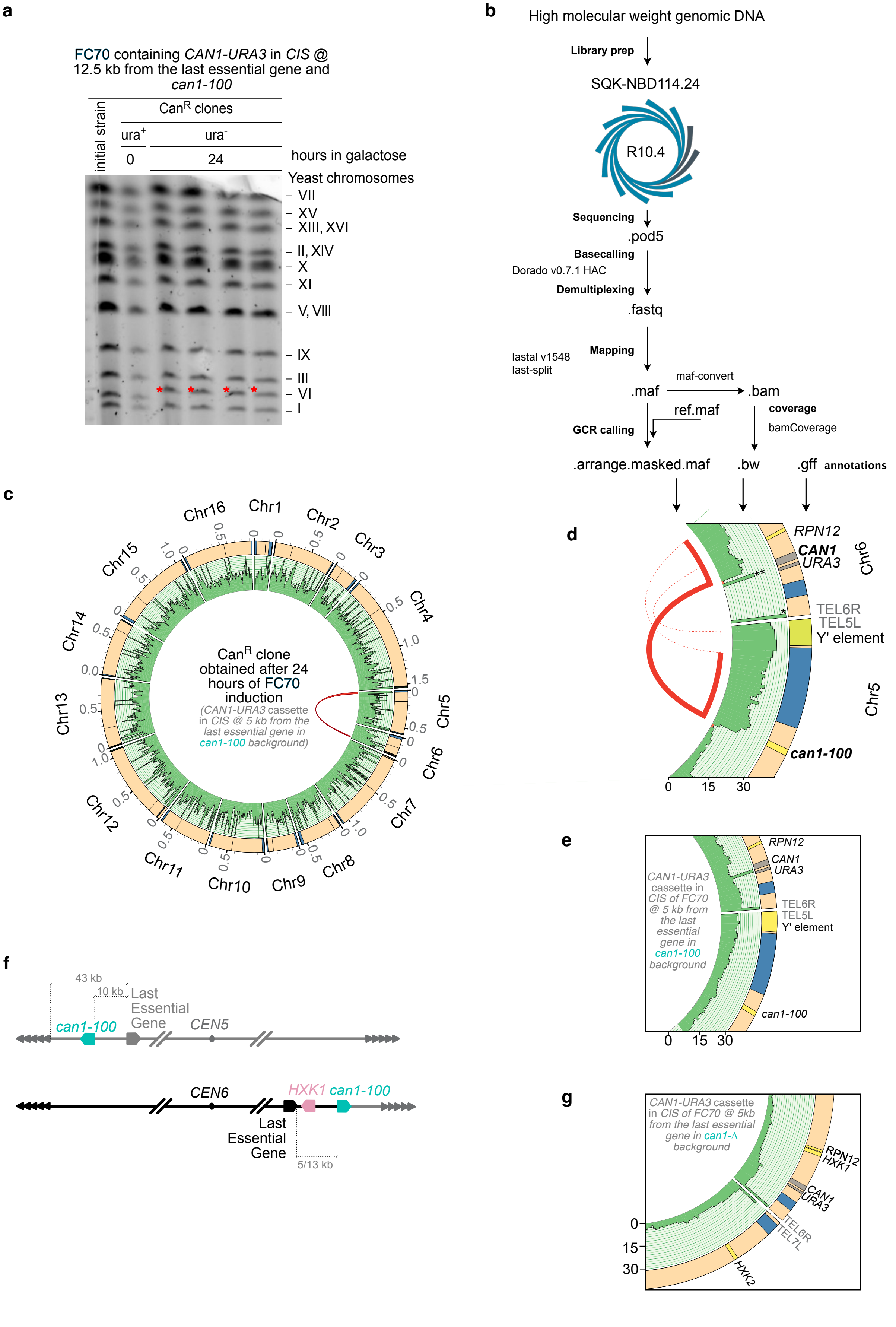
a
FC70 containing CAN1-URA3 in CIS @ 12.5 kb from the last essential gene and can1-100
initial strain
CanR clones
ura+
ura-
0
24
hours in galactose
Yeast chromosomes
VII
XV
XIII, XVI
II, XIV
X
XI
V, VIII
IX
III
VI
I
b
High molecular weight genomic DNA
Library prep
SQK-NBD114.24
R10.4
Sequencing
.pod5
Basecalling
Dorado v0.7.1 HAC
Demultiplexing
.fastq
Mapping
lastal v1548
last-split
.maf
maf-convert
.bam
ref.maf
GCR calling
coverage
bamCoverage
.arrange.masked.maf
.bw
.gff
annotations
c
Chr1
Chr2
Chr3
Chr4
Chr5
Chr6
Chr7
Chr8
Chr9
Chr10
Chr11
Chr12
Chr13
Chr14
Chr15
Chr16
CanR clone obtained after 24 hours of FC70 induction
(CAN1-URA3 cassette in CIS @ 5 kb from the last essential gene in can1-100 background)
d
RPN12
CAN1
URA3
Chr6
TEL6R
TEL5L
Y' element
Chr5
can1-100
0
15
30
e
CAN1-URA3 cassette in CIS of FC70 @ 5 kb from the last essential gene in can1-100 background
RPN12
CAN1
URA3
TEL6R
TEL5L
Y' element
can1-100
0
15
30
f
43 kb
10 kb
Last Essential Gene
can1-100
CEN5
CEN6
HXK1
can1-100
Last Essential Gene
5/13 kb
g
CAN1-URA3 cassette in CIS of FC70 @ 5kb from the last essential gene in can1-Δ background
RPN12
HXK1
CAN1
URA3
TEL6R
TEL7L
HXK2
0
15
30

**h**

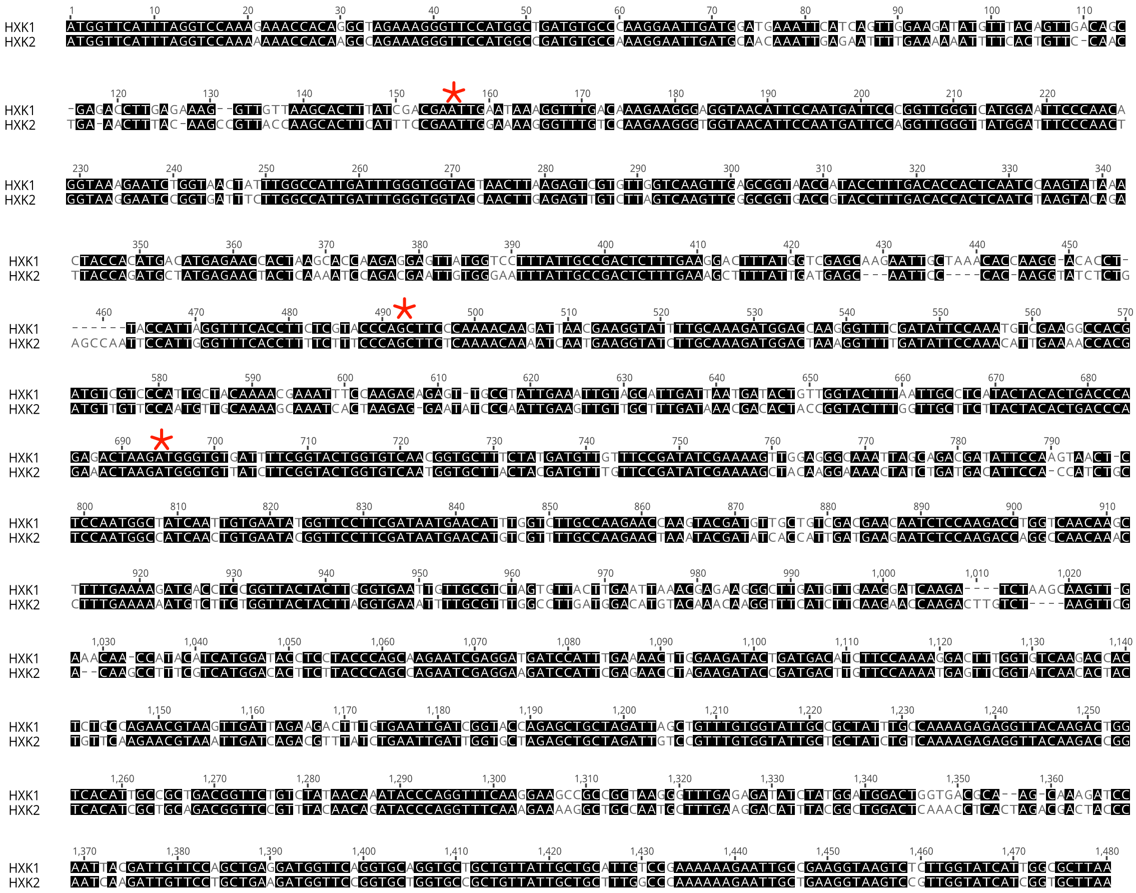

***Supplementary Fig. 7 (Related to Figure 7). Genomic instability at the shortest telomere in the cell enables senescence escape.***

***a,*** *Pulse-Field Gel Electrophoresis of indicated strains (derived from initial strain yT1696, grown for indicated time in galactose. Red asterisk indicates a change in chromosome 6 mobility. Chromosomes 4 and 12 run upper and were excluded from the picture.*

***b,*** *Pipeline for generating the CIRCOS based genome assembly.*

***c,*** *CIRCOS plot showing the genome of a Can$^R$ 5-FOA$^R$ ura$^-$ colony classified in this study as GCR, obtained after 24 h of Cas9 induction in the absence of telomerase of cells harboring a FC70 telomere, the CAN1-URA3 cassette at 5 kb from the last essential gene on chromosome 6R and in a can1-100 backgound (initial strain: yT1698, Supplementary Fig. 6b-c, f). Legends as in Fig. 7a.*

***d,*** *CIRCOS plot detail of c showing* loci *involved in the breakpoint region, including junction* CAN1-can1-100 *in bold. Chromosome 5 and 6 were tilted to achieve juxtapositition of* TEL5L *and* TEL6R *in the scheme. The solid red lines represents the longest splitted read. Dashed red lines indicate the extremities of splitted reads. (*) Reads containing exclusively TG$_{1-3}$, which map telomeric regions indiscriminately. (**) Reads mapping a region of the* CAN1-URA3 *cassette (in grey) containing a short plasmid-derived sequence (60 bp), also present at the* AFB2-TRP1 *locus.*

***e,*** *CIRCOS plot of the genome of control strain yT1698 zoomed into relevant region implicated in the rearangement showed in d.*

***f,*** *proposed model for the chromosomal rearrangement involving chromosomes 6 and 5 of Can$^R$.*

***g,*** *CIRCOS plot of genomes of control strain yT1760 zoomed into relevant regions implicated in the rearangement showed Fig. 7.*

***h,*** *Alignment of HXK1 and HXK2 present in yT1760. Homologous sequences are highlighted in black. Recombination regions characterized by ONT from three independent Can$^R$ mutants are indicated with red stars.*

### Supplementary Table 1: Yeast strains used in this work

| Name | Comment | Figure | Genotype |
|---|---|---|---|
| G49 | Obtained from M. Lisby | | MATα ura3-1 leu2-3,112 his3-11,15 can1-100 |
| yT992 | | | MATα ura3-1 leu2-3,112 his3-11,15 can1-100 tlc1::HIS3MX6-$P_{Teto2}$-TLC1 |
| yT1487 | yT992 transformed with Eco RV-digested YIp204-PADH1-AFB2 | S1,S4 | MATα ura3-1 trp1-1::AFB2-TRP1 leu2-3,112 his3-11,15 can1-100 tlc1::HIS3MX6-$P_{Teto2}$-TLC1 |
| yT1529 | yT1487/pT182 | S1,S4 | MATα ura3-1 trp1-1::AFB2-TRP1 leu2-3,112 his3-11,15 can1-100 tlc1::HIS3MX6-$P_{Teto2}$-TLC1 TEL6R::FC0 |
| yT1503 | yT1487/pT177 | 1,4,S1 | MATα ura3-1 trp1-1::AFB2-TRP1 leu2-3,112 his3-11,15 can1-100 tlc1::HIS3MX6-$P_{Teto2}$-TLC1 TEL6R::FC20 |
| yT1504 | yT1487/pT176 | 1-2,4,S1 | MATα ura3-1 trp1-1::AFB2-TRP1 leu2-3,112 his3-11,15 can1-100 tlc1::HIS3MX6-$P_{Teto2}$-TLC1 TEL6R::FC30 |
| yT1505 yT1527 | yT1487/pT175 | 1-2,4,S1 | MATα ura3-1 trp1-1::AFB2-TRP1 leu2-3,112 his3-11,15 can1-100 tlc1::HIS3MX6-$P_{Teto2}$-TLC1 TEL6R::FC40 |
| yT1506 | yT1487/pT174 | 1-2,4,S1 | MATα ura3-1 trp1-1::AFB2-TRP1 leu2-3,112 his3-11,15 can1-100 tlc1::HIS3MX6-$P_{Teto2}$-TLC1 TEL6R::FC50 |
| yT1576 | yT1507 backcrossed with G49 | S6 | MATa ura3-1 trp1-1::AFB2-TRP1 leu2-3,112 his3-11,15 can1-100 tlc1::HIS3MX6-$P_{Teto2}$-TLC1 |
| yT1597 | yT1487/pT187 | 1-4, S1-3 | MAT α ura3-1 trp1-1::AFB2-TRP1 leu2-3,112 his3-11,15 can1-100 tlc1::HIS3MX6-$P_{Teto2}$-TLC1 TEL6R::FC70 |
| yT1612 | yT1597 backcrossed with G49 | S6 | MATa ura3-1 trp1-1::AFB2-TRP1 leu2-3,112 his3-11,15 can1-100 tlc1::HIS3MX6-$P_{Teto2}$-TLC1 TEL6R::FC70 |
| yT1690 | CAN1-URA3 in trans FC70 | 6 | MATa ura3-1 trp1-1::AFB2-TRP1 leu2-3,112 his3-11,15 can1-100::CAN1-URA3 tlc1::HIS3MX6-$P_{Teto2}$-TLC1 TEL6R::FC70 |
| yT1694 | CAN1-URA3 in trans | 6 | MATa ura3-1 trp1-1::AFB2-TRP1 leu2-3,112 his3-11,15 can1-100::CAN1-URA3 tlc1::HIS3MX6-$P_{Teto2}$-TLC1 |
| yT1696 | CAN1-URA3 in cis FC70 @12.5kb of last essential gene | S6 | MATa ura3-1 trp1-1::AFB2-TRP1 leu2-3,112 his3-11,15 can1-100 tlc1::HIS3MX6-$P_{Teto2}$-TLC1 CHR06:266663::CAN1-URA3 TEL6R::FC70 |
| yT1698 | CAN1-URA3 in cis @ 5 kb of last essential gene | S6-7 | MATa ura3-1 trp1-1::AFB2-TRP1 leu2-3,112 his3-11,15 can1-100 tlc1::HIS3MX6-$P_{Teto2}$-TLC1 CHR06:258991::CAN1-URA3 TEL6R::FC70 |
| yT1710 | CAN1-URA3 in cis @12.5kb of last essential gene | S6 | MATa ura3-1 trp1-1::AFB2-TRP1 leu2-3,112 his3-11,15 can1-100 tlc1::HIS3MX6-$P_{Teto2}$-TLC1 CHR06:266663::CAN1-URA3 |
| yT1712 | CAN1-URA3 in cis @ 5 kb of last essential gene | S6 | MATa ura3-1 trp1-1::AFB2-TRP1 leu2-3,112 his3-11,15 can1-100 tlc1::HIS3MX6-$P_{Teto2}$-TLC1 CHR06:258991::CAN1-URA3 |
| yT1759 | CAN1-URA3 in cis @ 5 kb of last essential gene | 6 | MATa ura3-1 trp1-1::AFB2-TRP1 leu2-3,112 his3-11,15 can1-Δ tlc1::HIS3MX6-$P_{Teto2}$-TLC1 CHR06:258991::CAN1-URA3 |
| yT1760 | CAN1-URA3 in cis @ 5 kb of last essential gene | 6, S7 | MATa ura3-1 trp1-1::AFB2-TRP1 leu2-3,112 his3-11,15 can1-Δ tlc1::HIS3MX6-$P_{Teto2}$-TLC1 CHR06:258991::CAN1-URA3 TEL6R::FC70 |
| yT1869 | = yT1760 exo1-Δ | 6 | MATa ura3-1 trp1-1::AFB2-TRP1 leu2-3,112 his3-11,15 can1-Δ tlc1::HIS3MX6-$P_{Teto2}$-TLC1 CHR06:258991::CAN1-URA3 TEL6R::FC70 exo1-Δ |
| yT1870 | = yT1760 pol32-Δ | 6 | MATa ura3-1 trp1-1::AFB2-TRP1 leu2-3,112 his3-11,15 can1-Δ tlc1::HIS3MX6-$P_{Teto2}$-TLC1 CHR06:258991::CAN1-URA3 TEL6R::FC70 pol32-Δ |
| yT1871 | = yT1760 rad1-Δ | 6 | MATa ura3-1 trp1-1::AFB2-TRP1 leu2-3,112 his3-11,15 can1-Δ tlc1::HIS3MX6-$P_{Teto2}$-TLC1 CHR06:258991::CAN1-URA3 TEL6R::FC70 rad1-Δ |
| MCY813 | | 5 | tlc1-Δ::HIS3/TLC1 leu2-3,112/LEU2-$P_{RNR3}$-TLC1 ura3::caURA3-$P_{RPL24A}$-FHA1-mCherry/ura3::caURA3-$P_{RPL24A}$-FHA1-mCherry hta2::HTA2-ECFP-KANMX6/hta2::HTA2-ECFP-KANMX6 can1-100/can1-100 his3-11,15/his3-11,15 trp1-1/trp1-1 RAD5/RAD5 |

### Supplementary Table 2: Bacteria strains used in this work

| Name | Figure | Genotype |
|---|---|---|
| NEB® Stable Competent *E. coli* (High Efficiency) | 3, S3 | *F' proA+B+ lacIq Δ(lacZ)M15 zzf::Tn10 (TetR)/Δ(ara-leu) 7697 araD139 fhuA ΔlacX74 galK16 galE15 e14- Φ80dlacZΔM15 recA1 relA1 endA1 nupG rpsL (StrR) rph spoT1 Δ(mrr-hsdRMS-mcrBC)* |
| NEB® 10-beta Competent *E. coli* (High Efficiency) | | *Δ(ara-leu) 7697 araD139 fhuA ΔlacX74 galK16 galE15 e14- ϕ80dlacZΔM15 recA1 relA1 endA1 nupG rpsL (Str$^R$) rph spoT1 Δ(mrr-hsdRMS-mcrBC)* |

### Supplementary Table 3: Plasmids used in this work

| Name | Description | Source |
|---|---|---|
| pUC18 | Empty backbone. pUC cloning vector | [1] |
| bRA77 | Empty backbone. $P_{Gal1}$-*Cas9-AID-3myc*. gRNA can be cloned into BplI sites. Gift of J. Haber | [2] |
| bRA90 | Encodes $P_{PGK1}$-*Cas9-3xFlag*. Gift of J. Haber | [2] |
| pRS305 | Yeast integrative vector with a LEU2 marker. | [3] |
| pRS306 | Yeast integrative vector with a URA3 marker. | [3] |
| pHis-AID*-9myc | C-terminal AID*-9myc degron cassette with HIS3MX marker | [4] |
| Ylp204-PADH1-AFB2 | Expression of *PADH1-AFB2* | [4] |
| pT26 | pURTel[5] derivative to modify *TEL6R* | [6] |
| pT63 | Encodes $P_{PGK1}$-*Cas9-3xFlag;* sgRNA targets subtelomere 6R/*TEL6R* junction | This work |
| pT117 | Encodes $P_{Gal1}$-*Cas9-AID-3myc* in a replicative plasmid | This work |
| pT122 | Encodes $P_{Gal1}$-*Cas9-3xFlag* in an integrative plasmid | This work |
| pT126 | Encodes $P_{Gal1}$-*Cas9-AID-3myc;* sgRNA targets FC | This work |
| pT182 | For FC0 construction, contains fragment composed of subtelomere 6R followed by 5’-AAGCTTTTAGCATACGTGTGCGTACGCTCGAG**CCTGGGATCGCAGTGGTGAGTAA***GAGCTC*GTGGGTGTGG…-3’[1] | This work |
| pT0177 | For FC20 construction, contains fragment composed of subtelomere 6R followed by 5’-AAGCTTTTAGCATACGTGTGCGTACGCTCGAG**GTGGTGTGGGTGTGGTGTGT****CCTGGGATCGCAGTGGTGAGTAA***GAGCTC*GTGGGTGTGG…-3’[1] | This work |
| pT176 | For FC30 construction, contains fragment composed of subtelomere 6R followed by 5’-AAGCTTTTAGCATACGTGTGCGTACGCTCGAG**GTGGTGTGGGTGTGGTGTGTGGGTGTGGTG****CCTGGGATCGCAGTGGTGAGTAA***GAGCTC*GTGGGTGTGG…-3’[1] | This work |
| pT175 | For FC40 construction, contains fragment composed of subtelomere 6R followed by 5’-AAGCTTTTAGCATACGTGTGCGTACGCTCGAG**GTGGTGTGGGTGTGGTGTGTGGGTGTGGTGTGTGGGTGTG****CCTGGGATCGCAGTGGTGAGTAA***GAGCTC*GTGGGTGTGG…-3’[1] | This work |
| pT174 | For FC50 construction, contains fragment composed of subtelomere 6R followed by 5’-AAGCTTTTAGCATACGTGTGCGTACGCTCGAG**GTGGTGTGGGTGTGGTGTGTGGGTGTGGTGTGTGGGTGTGGTGTGGGTGT****CCTGGGATCGCAGTGGTGAGTAA***GAGCTC*GTGGGTGTGG…-3’[1] | This work |
| pT187 | For FC70 construction, contains fragment composed of subtelomere 6R followed by 5’-AAGCTTTTAGCATACGTGTGCGTACGCTCGAG**GTGGTGTGGGTGTGGTGTGTGGGTGTGGTGTGTGGGTGTGGTGTGGGTGTGTGGGTGTGTGGGTGTGGT****CCTGGGATCGCAGTGGTGAGTAA***GAGCTC*GTGGGTGTGG…-3’[1] | This work |
| pT188 | For FC100 construction, contains fragment composed of subtelomere 6R followed by 5’-AAGCTTTTAGCATACGTGTGCGTACGCTCGAG**GGTGGTGTGGGTGTGGTGTGTGGGTGTGGTGTGTGGGTGTGGTGTGGGTGTGTGGGTGTGTGGGTGTGGTGTGTGTGTGTGGGTGTGGGTGTGGGTGTGG****CCTGGGATCGCAGTGGTGAGTAA***GAGCTC*GTGGGTGTGG…-3’[1] | This work |
| pIK2 | $P_{RNR3}$-*TLC1* inserted into pRS305 | This work |
| pT194 | Encodes $P_{PGK1}$-*Cas9-Flag*. sgRNA targets *CAN1* locus | This work |
| pT196 | *CAN1* locus and flanking region (ChrV 31457-34131) cloned into pRS306. Plasmid for CAN1-URA3 cassette amplification. | This work |

[1] bold blue: telomeric repeats left at *FinalCut* telomere; **bold black: PAM and target sequence for Cas9 (reverse complement)**; *italics:* Sac *I site*; blue: terminal telomeric repeats.

### Supplementary Table 4: Oligos used in this work

| Name | 5′-3′ sequence |
|---|---|
| oT0883 | ACGTTTAGCTGAGTTTAACGGTG |
| oT1310 | GCGGATCCGGGGGGGGGG |
| oT1324 | AACAAAGTTCGGGATTCAGCTCG |
| oT1450 | GCCATATCAATATGCTAGTGGTTTT |
| oT1451 | CACTAGCATATTGATATGGCGATCA |
| oT1550 | TTACTCACCACTGCGATCCCGTTTT |
| oT1551 | GGGATCGCAGTGGTGAGTAAGATCA |
| oT1572 | ATGCACTAGTTGCACTAGGCG |
| oT1666 | CTCCAGATTTATCAGCAATAAAC |
| oT1667 | GTTTATTGCTGATAAATCTGGAG |
| oT1713 | AGCTTTTAGCATACGTGTGCGTACGC |
| oT1714 | TCGAGCGTACGCACACGTATGCTAAA |
| oT1741 | GACTACAAAGACCATGACGGTGATTATCGTACGCTGCAGGTCGAC |
| oT1742 | GTCGACCTGCAGCGTACGATAATCACCGTCATGGTCTTTGTAGTC |
| oT1749 | GCCTGGATGACACGTAAATCAG |
| oT1750 | CTGATTTACGTGTCATCCAGGC |
| oT1806 | CTGTTTAGCTTGCCTCGTCCCTGATAATCTCTTCTCGAGTCATGT |
| oT1808 | TCTCAATTGGGTGGTGATGAAGGGCGTACGCTGCAGGTCGAC |
| oT1829 | CCTGTAGCATCGATAGCAGC |
| oT1830 | GTTCGTTCCAATGTCAAGTTC |
| oT1831 | CAAATGGTGAGACTGGAGAG |
| oT1832 | GAAATCACGCCCCTTTGTCCC |
| oT1833 | TCACTCAAAGGCGGTAATAC |
| oT1834 | CCTTTTGCTCACATGTTCTTT |
| oT1835 | GAGTGCACCATATCGACTAC |
| oT1836 | CCTTCTTGAACCATTTCCCA |
| oT1881 | TTACTCACCACTGCGATCCC |
| oT1894 | CGTGGATGATGTGGTCTCTACAG |
| oT1959 | GGCTGGACTACTTTCTGGAATAGCG |
| oT1960 | ACTGAGTTCGGATCACTACACACGG |
| oT1961 | GATCCTAACGAGTGGATGCACAG |
| oT1962 | GACCCAGTCCTCATTTCCATC |
| oT1963 | GATGGTGGGGCAATTTCGAGAG |
| oT1964 | CATGGCCATTCCTCAGGATCTTC |
| oT1965 | CGATACCACGGCATTGATAAGC |
| oT1966 | GGCACACTCGTACCATAAAACAG |
| oT2028 | ATAGACCGAGATAGGGTTGAGTG |
| oT2029 | GTGGACTCTTGTTCCAAACTGG |
| oT2032 | CACACCCACACACCACACCCA |
| FW_RNR3pr+HindIII | ATCCTGAAGCTTAAGAGAAGGTAACAAGCACAT |
| REV_RNR3pr+BamHI | CAGGATGGATCCTTGTGTGGGAGTATTTGATTTATTG |
| FW_TLC1+BamHI | ATCCTGGGATCCTGGTTTGAGAATAAAACTAGAGAGG |
| REV_TLC1+XbaI | CAGGATTCTAGAAAGAAGAAGCCATTTGGTGG |
| oT2057 | ATGCAAGCTTTGAAATACCGGTTTT |
| oT2058 | CGGTATTTCAAAGCTTGCATGATCA |
| oT2082 | GGCGCGTCAGCGGGTGTTGGTATATCTTTAACAGATTCCAAAC |
| oT2083 | GTTTGGAATCTGTTAAAGATATACCAACACCCGCTGACGCGCC |
| oT2084 | GTAGTAAGCGCAACATACACCGGGTGTCGGGGCTGGC |
| oT2085 | GCCAGCCCCGACACCCGGTGTATGTTGCGCTTACTAC |
| oT2098 | TCCGGCACCAAGAATAGAGT |
| oT2126 | GATTTTTTAGTTCGATTTCATTCAGTAGAGAAAGGCATATCAATATGACATGTATATCTTTAACAGATTCCAAAC |
| oT2127 | TCTCTATTACGGTATTCTTTTTATTGCTAGCATTTAGCCGCGGAGTGAGATCAGTTTTGCTGGCCGCATCTTC |
| oT2130 | GAAAATGCTGGTGTGAATGTGAATGACGATAGACGGACTGATGCACTTTTCCATTATATCTTTAACAGATTCCAAAC |
| oT2131 | AGTCAAATTTTCAATCATGCTTCTCCCAATCTTGTAAGTAATGTTATCGTACAGTTTTGCTGGCCGCATCTTC |
| oT2132 | TTTCTGTTGGTGCTGATATTGCNNNNNNNNNNACGTTTAGCTGAGTTTAACGGTG |
| oT2139 | CCCAAAACAGTGCACCCAAG |
| oT2158 | ACTTGCCTGTCGCTCTATCTTCGGGGGGGGGGG |
| oT2162 | GAAGATAGAGCGACAGGCAAGTGCATGCAAGCTTGGCACTGG |
| oT2163 | GCAATATCAGCACCAACAGAAAGAATTCGTAATCATGGTCATAGCTGT |
| oT2189 | ATCGAACTGTGCGTGGAGAA |
| oT2190 | TTCTTGGAGGCAGGGGATTG |
| oT2199 | AGGCCACAGAACCGTATTCA |
| oT2276 | TCAATCACTTACTGGCAAGTGCGTATAAATTAAACCTATTTCTTTATCATCATATTTACTTAAACCATTAAAGAATA<br>TCTCGACCAGAATCTAACAGATATACATGTTCCGATAATGTCT |
| oT2357 | ACTTGCCTGTCGCTCTATCTTCGGGGGGGGGGGGGGGGGG |

### Supplementary Table 5: Composition of buffers used in Southern blots

| | Hybond XL RPN303B GE Healthcare Amersham | Hybond N+RPN303B Amersham Hybond N+RPN303B |
|---|---|---|
| *Hybridization buffer* | Church buffer: 500 mM sodium phosphate pH 7.4, 1 mM EDTA; 7% SDS, 1% BSA | Modified Church and Gilbert buffer: phosphate buffer 0,5 M pH 7,2; SDS 7%; EDTA 10 mM |
| *Low stringency wash* | 2× SSC and 0.1% SDS – 2 brief washes at RT | 2x SSC, 0,1% SDS – one brief wash at RT; two washes 5 min 65°C |
| *Medium stringency wash* | 1× SSC and 0.1% SDS – 1 wash of 15 min at 65°C | 1x SSC, 0,1 %SDS – two washes 10 min each 65°C |
| *Medium stringency wash* | 0,5× SSC and 0.1% SDS – 1 wash of 15 min at 65°C | |
| *High stringency wash* | 0,1× SSC and 0.1% SDS– 1-2 washes of 10 min at 65°C | 0,1× SSC and 0.1% SDS – 4 washes 5 min at 65°C |

### Supplementary Table 6: Average ± s.e.m. (n=3) of fraction of uncut and cut bands in southern as in Fig.1d-e.

| *FinalCut* | Hours in galactose | uncut | cut |
|---|---|---|---|
| *FC20* | 0 | 1 ± 0 | 0 ± 0 |
| | 3 | 0,518 ± 0,169 | 0,091 ± 0,018 |
| | 6 | 0,216 ± 0,101 | 0,043 ± 0,017 |
| | 9 | 0,1 ± 0,03 | 0 ± 0 |
| *FC30* | 0 | 1 ± 0 | 0 ± 0 |
| | 3 | 0,575 ± 0,074 | 0,24 ± 0,02 |
| | 6 | 0,191 ± 0,022 | 0,167 ± 0,009 |
| | 9 | 0,102 ± 0,026 | 0,207 ± 0,04 |
| *FC40* | 0 | 1 ± 0 | 0 ± 0 |
| | 3 | 0,839 ± 0,076 | 0,15 ± 0,033 |
| | 6 | 0,369 ± 0,074 | 0,28 ± 0,054 |
| | 9 | 0,177 ± 0,051 | 0,32 ± 0,052 |
| *FC50* | 0 | 1 ± 0 | 0 ± 0 |
| | 3 | 0,427 ± 0,05 | 0,374 ± 0,058 |
| | 6 | 0,116 ± 0,031 | 0,513 ± 0,077 |
| | 9 | 0,078 ± 0,008 | 0,519 ± 0,033 |
| *FC70* | 0 | 0,991 ± 0,007 | 0,009 ± 0,007 |
| | 3 | 0,554 ± 0,064 | 0,459 ± 0,055 |
| | 6 | 0,21 ± 0,094 | 0,805 ± 0,037 |
| | 9 | 0,039 ± 0,021 | 0,967 ± 0,187 |

### Supplementary Table 7: Fraction of cells in which the shortest telomere is the *FinalCut* telomere, computed from simulations of 50,000 lineages

| *FinalCut* | Fraction of lineages in which the shortest telomere after the cut is *FinalCut* |
|---|---|
| *FC20* | 1 |
| *FC30* | 1 |
| *FC40* | 1 |
| *FC50* | 1 |
| *FC70* | 0.999896773127980 |

### Supplementary Table 8: Parameters fitted on the microfluidic data and used for the simulations

| parameter | Value for the general model[7] | *FinalCut* (this work) |
|---|---|---|
| $a_{nta}$ | 0.02 | 0.02 |
| $b_{nta}$ | 0.44 | 0.44 |
| $a_{sen,A}$ | 0.19 | 0,17 |
| $b_{sen,A}$ | 0.73 | 0.65 |
| $a_{sen,B}$ | 0 | 0 |
| $b_{sen,B}$ | 0.12 | 0.12 |
| $l_{min,A}$ | 27 | 36 |
| $l_{min,B}$ | 0 | 0 |
| $l_{trans}$ | 0 | 0 |
| $l_0$ | 40 | 40 |

$l_1$ 58 58

### Supplementary Table 9: Set of strains to study genomic instability in this work

| position of *CAN1-URA3* cassette | *FC* status | *CAN1* @ chr 5L | Other mutations | strain name | Figure |
|---|---|---|---|---|---|
| In *trans* @ subtelomere 5L (@ *can1-100* locus) | *FC70* | *can1-100* | | yT1690 | 6a,c,e |
| In *trans* @ subtelomere 5L (@ *can1-100* locus) | *noFC* | *can1-100* | | yT1694 | 6a,c,e |
| In *cis* @12.5 kb from last essential gene on 6R arm | *FC70* | *can1-100* | | yT1696 | S6a,e,S7a |
| In *cis* @12.5 kb from last essential gene on 6R arm | *noFC* | *can1-100* | | yT1710 | S6e |
| In *cis* @ 5 kb from last essential gene on 6R arm | *FC70* | *can1-100* | | yT1698 | S6b,f,S7c-f |
| In *cis* @ 5 kb from last essential gene on 6R arm | *noFC* | *can1-100* | | yT1712 | S6f |
| In *cis* @ 5 kb from last essential gene on 6R arm | *FC70* | *can1-Δ* | | yT1760 | 6b,d,f,7a-b,S7g-h |
| In *cis* @ 5 kb from last essential gene on 6R arm | *noFC* | *can1-Δ* | | yT1759 | 6d |
| In *cis* @ 5 kb from last essential gene on 6R arm | *FC70* | *can1-Δ* | *pol32-Δ* | yT1870 | 6f |
| In *cis* @ 5 kb from last essential gene on 6R arm | *FC70* | *can1-Δ* | *exo1-Δ* | yT1869 | 6f |
| In *cis* @ 5 kb from last essential gene on 6R arm | *FC70* | *can1-Δ* | *rad1-Δ* | yT1871 | 6f |

### Supplementary Table 10: Results of fluctuation analyses applying canavanine selection

| *CAN1-URA3* cassette position | Time (h) | Relevant genotype | *CAN1* Mutation rate x $10^{-6}$ [95% Confidence interval] | n[1] | Mutation type distribution[a]: Point mutation | GCR | Others | n[2] |
|---|---|---|---|---|---|---|---|---|
| 10 kb from the LEG[b] on arm 5L | 0 | *can1-100::CAN1-URA3 NoFC* | 0.1 [0.08-0.12] | 10 | 100 | 0 | 0 | 99 |
| | | *can1-100::CAN1-URA3 FC70* | 0.08 [0.07-0.10] | 16 | 100 | 0 | 0 | 68 |
| | 24 | *can1-100::CAN1-URA3 NoFC* | 0.08 [0.06-0.10] | 10 | 100 | 0 | 0 | 48 |
| | | *can1-100::CAN1-URA3 FC70* | 0.12 [0.09-0.16] | 13 | 96.25±5.3 | 3.75±5.3 | 0 | 48 |
| 12.5 kb from the LEG[b] on arm 6R | 0 | *can1-100 NoFC* | 0.33 [0.29-0.37] | 9 | 54.5±6.4 | 45.5±6.4 | 0 | 99 |
| | | *can1-100 FC70* | 0.66 [0.57-0.76] | 5[c] | 24.25±3.2 | 71.75±0.35 | 4±2.82 | 99 |
| | 24 | *can1-100 NoFC* | 0.37 [0.32-0.43] | 14 | 36.75±1.8 | 63.25±1.77 | 0 | 92 |
| | | *can1-100 FC70* | 8.26 [7.41-9.14] | 5[c] | 1±1.4 | 99±1,4 | 0 | 91 |
| 5 kb from the LEG[b] on arm 6R | 0 | *can1-100 NoFC* | 0.18 [0.16-0.20] | 15 | 76.7±9.9 | 20.9±13.34 | 2.4±4.22 | 129 |
| | | *can1-100 FC70* | 0.41 [0.37-0.44] | 15 | 30.7±15.3 | 65.3± 21.0 | 4±6.9 | 150 |
| | 24 | *can1-100 NoFC* | 0.16 [0.13-0.19] | 15 | 61.7±4.5 | 38.3±4.47 | 0 | 128 |
| | | *can1-100 FC70* | 6.96 [6.53-7.39] | 15 | 2±2 | 97.3±1.15 | 0.7±1.15 | 200 |
| | 0 | *can1-Δ NoFC* | 0.12 [0.11-0.14] | 19 | 100 | 0 | 0 | 239 |
| | | *can1-Δ FC70* | 0,19 [0.16-0.22] | 16 | 100 | 0 | 0 | 183 |
| | 24 | *can1-Δ NoFC* | 0.15 [0.12-0.18] | 18 | 99.6±0.7 | 0.4±0.7 | 0 | 203 |
| | | *can1-Δ FC70* | 0.68 [0.57-0.80] | 16 | 35.9±14.6 | 64.1±14.6 | | 313 |

n[1]: number of independent cultures
n[2]: number of Can[R] colonies analyzed
[a]Values are expressed in % ± SD over 2-3 independent experiments corresponding to n[1] cultures
[b]LEG: Last Essential Gene
[c]Experiment involving a single *FC70* transformant and experiment

### Supplementary Table 11: Results of fluctuation analyses applying canavanine and 5-FOA double selection

| *CAN1-URA3* cassette position | Relevant Genotype | Media | *CAN1-URA3* Mutation rate x$10^{-10}$ [95% Confidence interval] | n[1] |
|---|---|---|---|---|
| 10 kb from the LEG[b] on arm 5L (*TRANS*) | *can1-100::CAN1-URA3 FC70* | Raffinose | 1.05 [0.24-2.25] | 21 |
| | | Galactose | 2.11 [0.69-4.06] | 21 |
| 5 kb from the LEG[b] on arm 6R (*CIS*) | *can1-Δ FC70* | Raffinose | 58.31 [41.31-77.39] | 12 |
| | | Galactose | 1044.99 [904.58-1192.76] | 12 |
| | *can1-Δ pol32-Δ FC70* | Raffinose | 1.05 [0.19-2.37] | 17 |
| | | Galactose | 35.94 [21.65-52.77] | 17 |
| | *can1-Δ exo1-Δ FC70* | Raffinose | 35.89 [19.18-56.21] | 10 |
| | | Galactose | 933.92 [758.38-1122.63] | 10 |
| | *can1-Δ rad1-Δ FC70* | Raffinose | 136.69 [95.60-183.00] | 10 |
| | | Galactose | 1956.30 [1639.40-2293.50] | 10 |

n[1]: number of independent cultures
[b]LEG: Last Essential Gene

### Supplementary Table 12: Results of fluctuation analyses applying double selection – Ratios Gal/Raff

| | *TRANS* | *CIS* | | | |
|---|---|---|---|---|---|
| | *WT* | *WT* | *pol32* | *exo1* | *rad1* |
| ratio Gal/Raff | 2.01 | 17.92 | 34.23 | 26.02 | 14.31 |

### Supplementary Table 13: Results of fluctuation analyses applying double selection – Ratios mutant/WT

| | *pol32* | | *exo1* | | *rad1* | |
|---|---|---|---|---|---|---|
| | Raff | Gal | Raff | Gal | Raff | Gal |
| ratio mutant/WT | 0.02 | 0.03 | 0.62 | 0.89 | 2.34 | 1.87 |

### Supplementary Pipeline 1


```
<geneiousWorkflows>
  <XMLSerialisableRootElement name="TT_Clone&amp;Nanopore_Analysis1.1" author="teresatelott"
geneiousVersion="2023.2.1" uniqueId="d84ce14a-613a-490c-a277-fb89c20e7d5e" revisionNumber="15" description=""
bundledIconName="plugin">
    <workflowElement type="com.bioMATters.plugins.workflows.WorkflowElementForEach" />
    <workflowElement id="Geneious" exposeNoOptions="true" exposeAllOptions="false" suppressErrors="false"
showButtonForExposedGroup="false" groupNameForExposedOptions=""
type="com.bioMATters.plugins.workflows.DocumentOperationWorkflowElement">
      <Options>
        <option name="determineDirection">false</option>
        <option name="alignmentType">globalFreeEndGaps</option>
        <option name="nucleotideCostMATrix">70% similarity (IUB)</option>
        <option name="proteinCostMATrix">Blosum62</option>
        <option name="gapOpenPenalty">25.0</option>
        <option name="gapExtensionPenalty">5.0</option>
        <option name="refinementIterations">10</option>
        <option name="fastGuideTree">false</option>
        <option name="dontAlign">false</option>
      </Options>
      <optionToExpose optionName="determineDirection" label="" />
    </workflowElement>
    <workflowElement id="FilterOperation" exposeNoOptions="true" exposeAllOptions="false" suppressErrors="false"
showButtonForExposedGroup="false" groupNameForExposedOptions=""
type="com.bioMATters.plugins.workflows.DocumentOperationWorkflowElement">
      <Options>
        <option name="filterWhat">eachSequence</option>
        <option name="MATch">all</option>
        <multiOption name="filter">
          <value>
            <option name="field">percentage_similarity</option>
            <option name="condition">greater_than</option>
            <option name="value">95</option>
          </value>
        </multiOption>
      </Options>
      <optionToExpose optionName="filterWhat" label="" />
    </workflowElement>
    <workflowElement id="FilterOperation" exposeNoOptions="true" exposeAllOptions="false" suppressErrors="false"
showButtonForExposedGroup="false" groupNameForExposedOptions=""
type="com.bioMATters.plugins.workflows.DocumentOperationWorkflowElement">
      <Options>
        <option name="filterWhat">eachSequence</option>
```

```xml
          <option name="MATch">all</option>
          <multiOption name="filter">
            <value>
              <option name="field">percentage_identical</option>
              <option name="condition">greater_than</option>
              <option name="value">79.99999</option>
            </value>
          </multiOption>
        </Options>
        <optionToExpose optionName="filterWhat" label="" />
      </workflowElement>
      <workflowElement id="FilterOperation" exposeNoOptions="true" exposeAllOptions="false" suppressErrors="false"
showButtonForExposedGroup="false" groupNameForExposedOptions=""
type="com.bioMATters.plugins.workflows.DocumentOperationWorkflowElement">
        <Options>
          <option name="filterWhat">eachSequence</option>
          <option name="MATch">all</option>
          <multiOption name="filter">
            <value>
              <option name="field">meanCoverage</option>
              <option name="condition">greater_than</option>
              <option name="value">4.99999</option>
            </value>
          </multiOption>
        </Options>
        <optionToExpose optionName="filterWhat" label="" />
      </workflowElement>
      <workflowElement id="Generate_Consensus" exposeNoOptions="true" exposeAllOptions="false" suppressErrors="false"
showButtonForExposedGroup="false" groupNameForExposedOptions=""
type="com.bioMATters.plugins.workflows.DocumentOperationWorkflowElement">
        <Options>
          <option name="thresholdPercent">0</option>
          <option name="thresholdPercentNoQuality">65</option>
          <option name="noConsensusGaps">false</option>
          <option name="mapQuality">false</option>
          <option name="mapQualityMethod">mapSummed</option>
          <option name="noCoverageCharacterDeNovo">gap</option>
          <option name="noCoverageCharacterReference">gap</option>
          <option name="applyLowCoverageOrQualityCall">true</option>
          <option name="lowCoverageOrQualityCharacter">gap</option>
          <option name="coverageOrQuality">coverage</option>
          <option name="qualityThreshold">20</option>
          <option name="coverageThreshold">5</option>
          <option name="splitAroundQuestionMarks">false</option>
          <option name="noConsensusEndGaps">true</option>
          <option name="trimToReference">false</option>
          <option name="ignoreReadsMappedToMultipleLocations">false</option>
          <option name="removeGaps">true</option>
          <option name="appendText">true</option>
          <option name="textToAppend">consensus sequence</option>
          <option name="callWhenGapInBestStates" />
          <option name="callChroMATogramHeterozygotes">false</option>
          <option name="chroMATogramHeterozygotePercentage">50</option>
          <option name="howToStoreSequences">AskUser</option>
        </Options>
        <optionToExpose optionName="thresholdPercent" label="" />
      </workflowElement>
      <workflowElement type="com.bioMATters.plugins.workflows.WorkflowElementGroupSequences" />
      <workflowElement id="batchRename" exposeNoOptions="true" exposeAllOptions="false" suppressErrors="false"
showButtonForExposedGroup="false" groupNameForExposedOptions=""
type="com.bioMATters.plugins.workflows.DocumentOperationWorkflowElement">
        <Options>
          <option name="advancedCheckbox">false</option>
          <option name="customComponent1" />
```

```
        <childOption name="renameMethodChildOptions">
          <option name="selectedRenameMethod">renamableOption0</option>
          <childOption name="BatchRenameSubOptions0">
            <option name="attributeProcedure">nameProcedure</option>
            <option name="startBatchIndexOption">0</option>
            <option name="endBatchIndexInclusiveOption">2147483647</option>
            <option name="sequencesInSequenceListDocumentRenamableSequenceType">allSequencesInListOption</option>
          </childOption>
          <childOption name="BatchRenameSubOptions1">
            <option name="attributeProcedure">nameProcedure</option>
            <option name="followAndRenameReferences">true</option>
          </childOption>
          <childOption name="BatchRenameSubOptions2">
            <option name="selectedFieldComboOption">fieldOptioncache_name</option>
          </childOption>
        </childOption>
        <childOption name="basicSuperOptions">
          <option name="task">replaceWithRadioButton</option>
          <option name="r1">fieldOptiondocument_folder</option>
          <option name="rSep"> </option>
          <option name="r2">fieldOptioncache_name</option>
          <option name="rSep2" />
          <option name="r3">nothingSelectedOption</option>
          <option name="appendTextField" />
          <option name="appendPositionComboBox">endValue</option>
          <option name="removeHowMuchIntegerOption">1</option>
          <option name="removePositionComboBox">endValue</option>
        </childOption>
        <childOption name="advancedSuperOptions">
          <option name="replaceWhatTypeRadioOption">entireFieldOption</option>
          <option name="replaceWhatTextField" />
          <option name="ignoreCaseCheckbox">true</option>
          <option name="isRegexCheckbox">false</option>
          <option name="replaceWithTextField" />
          <option name="plusButton">Add Property...</option>
        </childOption>
      </Options>
      <optionToExpose optionName="advancedCheckbox" label="" />
    </workflowElement>
  </XMLSerialisableRootElement>
</geneiousWorkflows>
```

### Supplementary Pipeline 2

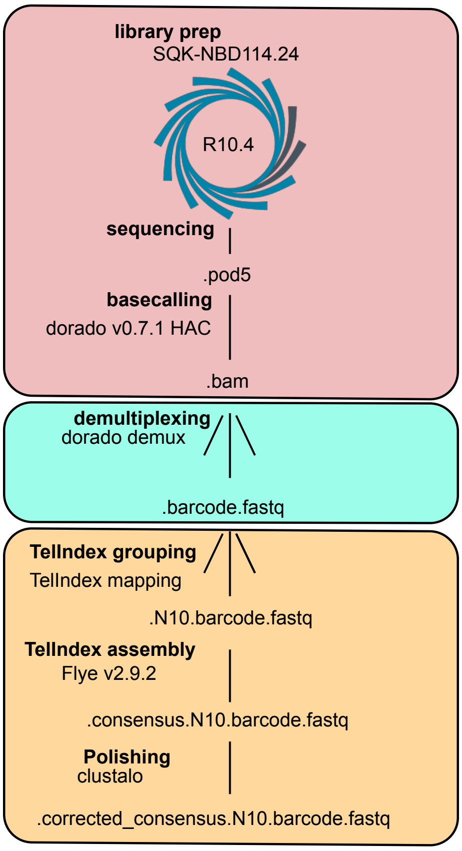


*Analysis pipeline for high throughput individual telomere sequencing. DNA from TeloPCRs is prepared according to the SQK-SBD114.24 library protocol on a R10.4 Nanopore flowcell. Raw signal is then basecalled and demultiplexed. Individual telomeres are grouped according to their TelIndex tag. Grouped reads are assembled and consensus telomeres are multialigned in order to correct potential sequencing/assembly errors.*
